# A Two-Stage Interpretable Framework for Predicting Plant-Derived Small RNA Targets on Human 3’UTRs

**DOI:** 10.64898/2026.06.16.732785

**Authors:** Le Qiao, Weizhong Li

## Abstract

Can plant-derived small RNAs target human mRNA 3’UTRs via complementary base pairing and produce experimentally detectable regulatory effects? This question concerns not only the fundamental feasibility of cross-kingdom RNA regulation but also the technological pathway for screening plant-derived active small nucleic acids. Existing miRNA target prediction tools are predominantly designed for endogenous miRNA-mRNA systems, exhibiting notable limitations when applied to cross-species small RNA inputs and small-sample wet-lab experimental adaptation. In this study, we developed a two-layer prediction framework, MetaLulu-AI. The first layer builds upon publicly available human miRNA-mRNA 3’UTR interaction data, utilizing XGBoost to learn foundational binding rules on human 3’UTRs based on 41 interpretable computational features, including seed region pairing types, local context sequence composition, site positioning, and RNA secondary structures. The second layer is tailored to the experimental system of plant-derived small RNAs and human target genes. It introduces 40 experimental samples using significant changes in endogenous protein expression as the regulatory standard (determined by Western blot or ELISA 48 hours post-transfection of small RNAs via Lipo3000). Using 52-dimensional computational features and the optimal transcript scores from the first layer as inputs, this layer employs TabPFN for experimental label adaptation.

The first-layer dataset consists of 38,752 training samples, 5,536 validation samples, and 11,073 testing samples (totaling 55,361), with a positive-to-negative sample ratio of approximately 1:5.4. On the randomly split test set, the model achieved an AUC of 0.9686, a recall of 0.8523, a precision of 0.8080, and an accuracy of 0.9452 (at a decision threshold of 0.4797). Group-based splitting revealed that the model maintains high discriminative power for unseen genes (AUC = 0.9541), though its generalization ability for completely unseen miRNAs decreases (AUC = 0.7390). For the 40 experimental samples in the second layer, the TabPFN model achieved an average AUC of 0.7406 ± 0.092 across ten repeated 70/30 random splits, outperforming the baseline of directly using the first-layer scores (0.3563 ± 0.149); the average AUC in a 5-fold cross-validation was 0.770 ± 0.177. SHAP analysis demonstrated a clear divergence in the discriminative basis of the two models: the first layer relies more heavily on the thermodynamics of the small RNA itself and the quality of canonical seed sites, whereas the second layer focuses more on the local UTR environment and statistical site features. Although the current second-layer results are constrained by sample size and gene coverage, this framework serves as a preliminary observation of the adaptation mechanism for cross-kingdom regulation experiments, and motivating future large-scale validation. Under stricter leave-one-gene-out and leave-one-small-RNA-out evaluation, the adapter exceeded the first-layer score baseline but only matched the majority-class baseline, underscoring that entity-level generalization is not yet established.

## 1 Introduction

### 1.1 Research Opportunities in Cross-Kingdom Small RNA Regulation

Small RNAs (sRNAs) participate in post-transcriptional gene expression regulation through complementary base pairing with target mRNAs. For endogenous miRNAs, factors such as the seed region, target site context, RNA secondary structure, and binding free energy all influence targeting recognition efficiency. Therefore, accurately predicting the binding relationship between sRNAs and mRNA 3’UTRs is a crucial prerequisite for candidate target screening and the design of subsequent functional experiments.

In recent years, whether plant-derived small RNAs can affect animal or human gene expression has gradually emerged as an important direction in cross-kingdom RNA regulation research. Existing studies suggest that there may be recognizable cross-kingdom interactions between plant sRNAs and animal/viral target sequences. For example, rice miR168a may enter mice through dietary intake and affect the expression of *LDLRAP1*; plant miR159 has been associated with the inhibition of breast cancer cell proliferation; and honeysuckle-derived miR2911 has been reported to target influenza virus-associated genes and influence viral replication.

However, the reproducibility and biological significance of these reported cross-kingdom regulatory events remain controversial. Dickinson et al. (2013) failed to replicate the regulatory effect of miR168a on *LDLRAP1* in independent experiments, while Witwer and Hirschi (2014) questioned whether the detection of plant miRNAs in plasma might be confounded by technical noise. Mechanistically, whether dietary exogenous small RNAs can reach target cells at sufficient concentrations and be incorporated into the RNA-induced silencing complex (RISC) still lacks adequate *in vivo* evidence.

In light of this, this paper does not attempt to argue for the universal existence of cross-kingdom regulation, but rather reframes the problem at a more tractable, operational level: given a plant-derived small RNA sequence and a human mRNA 3’UTR sequence, can we establish a reliable computational framework to provide an interpretable prioritization of their binding probabilities, thereby mitigating the guesswork and trial-and-error costs associated with subsequent wet-lab experiments?

### 1.2 Limitations of Existing Prediction Tools in Cross-Kingdom Scenarios

Classical computational tools such as TargetScan, miRanda, PITA, RNAhybrid, and miRDB have provided an essential foundation for miRNA target prediction. TargetScan focuses on seed region matching, site context, and evolutionary conservation; RNAhybrid and PITA are more concerned with binding free energy or site accessibility. miRDB is an approach that conceptually aligns more closely with our methodology; its core algorithm, MirTarget, does not rely solely on manual rules but is a machine learning model trained on known functional miRNA-mRNA interaction data. MirTarget translates sequence complementarity, seed region matching, target site context, and associated sequence features between the miRNA and candidate target sites into model inputs, outputting a prediction score from 0 to 100 to represent the likelihood of a candidate target being a functional miRNA target. The advantage of miRDB lies in its use of experimentally supported functional target information to train the model, avoiding a complete reliance on single seed rules; this shares similarity with our approach of learning foundational binding rules using publicly available human miRNA-mRNA data.

A fundamental constraint of miRDB lies in its training scope: the model learns from human endogenous miRNA-mRNA interaction data, operating under the implicit assumption that the input small RNAs are similar to human miRNAs in sequence composition and seed usage preferences. When the input switches to plant-derived small RNAs, this assumption no longer holds—plant miRNAs exhibit systematic differences from human miRNAs in length distribution, GC content, and seed region preferences. A more practical issue is that miRDB provides a single-stage target score learned from miRNA-target site features and does not explicitly incorporate downstream assay condition covariates such as cell lines, transfection efficiency, detection time windows, or Western blot/ELISA techniques. This motivated us to consider a two-stage strategy: first learning generalized binding rules, and then performing system-specific adaptation using small-sample experimental data.

Specifically, when plant-derived small RNAs cross-kingdom to target human mRNA 3’UTRs, the prediction task faces three primary challenges. First, there is usually a lack of directly utilizable evolutionary conservation of target sites between plants and animals. Second, binding patterns in cross-kingdom scenarios may not fully conform to classical canonical seed site rules. Third, wet-lab readouts are influenced by factors such as cellular background, transfection conditions, **reporter system construction methods**, and detection timeframes; a simple foundational binding probability score does not necessarily correspond directly to experimental positive or negative labels.

In recent years, several deep learning models, such as DeepMirTar, TargetNet, and Mimosa, have made progress in miRNA target prediction. However, the inherent “black box” nature of deep neural networks makes their decision mechanisms extremely difficult to interpret, presenting a barrier to use in scenarios where experimental design must be interpreted alongside biophysical mechanisms. For the task of screening plant-derived small RNA candidates, a model requires not only strong discriminatory power but also the ability to explain the basis of its predictions and adapt to specific experimental systems.

### 1.3 Current Work and Core Methodology

The core design philosophy of this study is to decompose the complex cross-kingdom target prediction task into two relatively independent subproblems. The first subproblem—“Does this small RNA possess the theoretical potential to bind the target 3’UTR at the sequence matching and thermodynamic levels?”—can be addressed using large volumes of annotated human miRNA-mRNA interaction data available in public databases. The second subproblem—“Can this binding propensity elicit a significant change in endogenous protein expression (e.g., as detected by Western blot or ELISA)?”—can only be calibrated using system-specific wet-lab experimental data. The proposed framework is built precisely around this decomposition strategy.

The first layer leverages public human miRNA-mRNA 3’UTR data to learn foundational binding rules, including interpretable signals such as seed region pairing, local context, site positioning, and RNA thermodynamics. The second layer employs wet-lab experimental data detailing changes in endogenous target protein expression following transfection with plant-derived small RNAs to experimentally adapt the first-layer output and the corresponding optimal transcript features. This design mitigates the overfitting risk associated with training complex models directly on a small dataset of 40 wet-lab experimental samples, while simultaneously avoiding the logical fallacy of simply equating the first layer’s “foundational binding probability” with the “final experimental phenotype.”

The main contributions of this paper are as follows:

1. **Construction of a feature engineering system:** We developed a 41-dimensional interpretable computational feature system tailored for small RNA-3’UTR binding recognition, which was utilized in the first layer to learn foundational binding rules.
2. **Exploration of the base model and generalization boundaries:** We trained an XGBoost base predictor using 55,361 publicly available human miRNA-mRNA 3’UTR samples. Its generalization boundaries and potential information leakage risks were evaluated through random splitting, gene-based group splitting, miRNA-based group splitting, and label-shuffling experiments.
3. **Small-sample experimental adaptation scheme:** We constructed a second-layer TabPFN experimental adaptation model tailored to 40 experimental samples assessing the regulation of endogenous protein expression by plant-derived small RNAs targeting human gene 3’UTRs. We subsequently evaluated its performance variations relative to the first-layer baseline through repeated random splits and 5-fold cross-validation.
4. **Attribution interpretation of the two-layer decision mechanism:** We employed SHAP analysis to interpret the two-layer model, analyzing the differences between the foundational binding rules and the experimental adaptation mechanisms.

## 2 Methods

### 2.1 Overall Framework

The proposed framework employs a two-layer prediction architecture (Figure 1).

**Figure 1:**
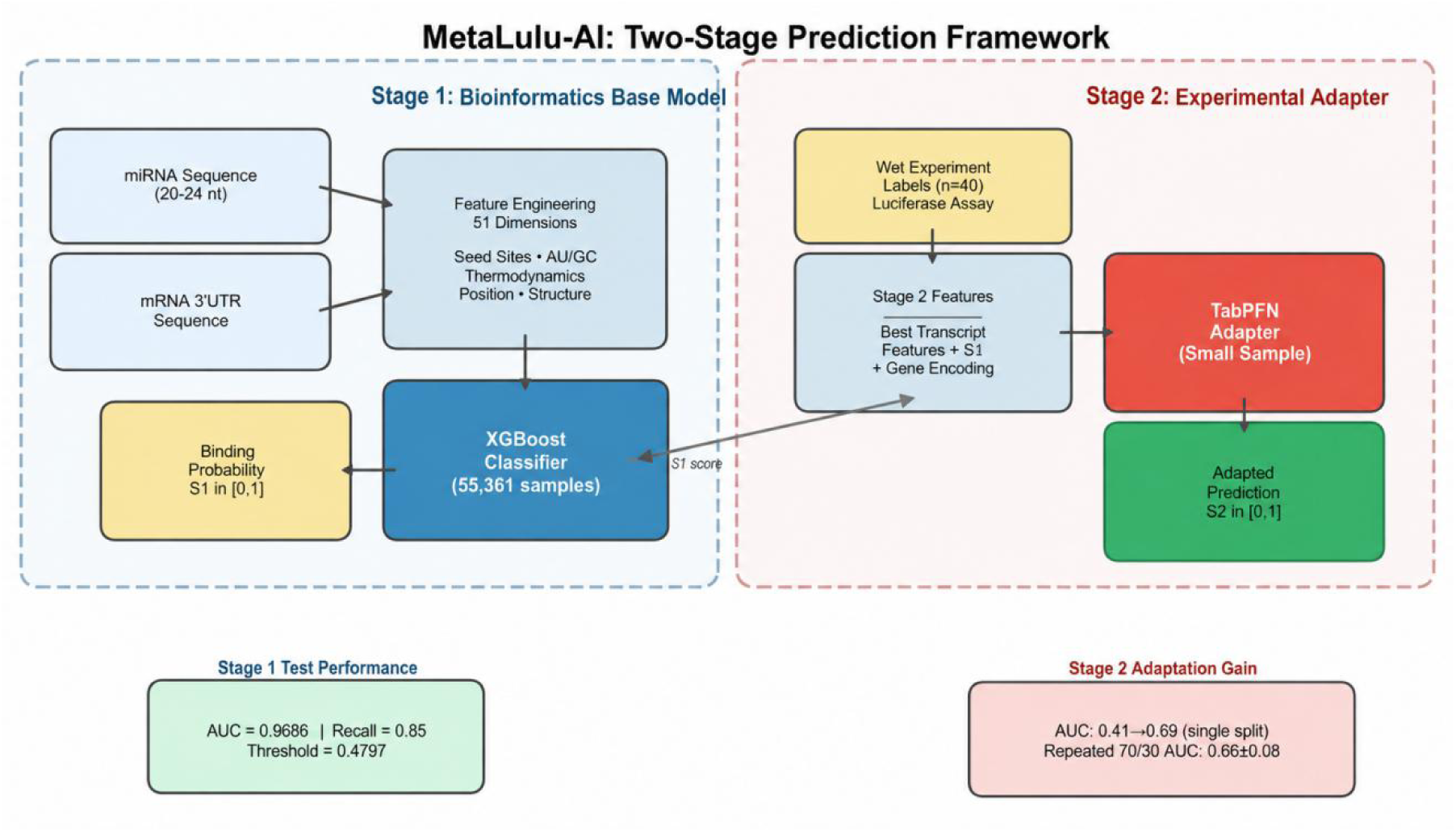
The Two-Layer Prediction Framework.

The first layer consists of a foundational XGBoost model. By taking small RNA sequences and target mRNA 3’UTR sequences as inputs, it extracts interpretable bioinformatics features and outputs a foundational binding probability score. Trained on annotated human miRNA-mRNA data, this layer is primarily utilized to learn the foundational binding rules on human 3’UTRs associated with seed region pairing, local context, and structural thermodynamics.

The second layer features a TabPFN model for experimental adaptation. The inputs are 52-dimensional final computational features, which include the optimal transcript score (best_score) generated by the first layer. The supervisory signals are derived from experimental labels based on changes in endogenous target protein expression following the transfection of plant-derived small RNAs. The second layer explicitly excludes identifier fields such as target gene names, miRNA names, sample IDs, or experimental labels.

The first and second layers have distinct modeling objectives. The first layer aims to provide foundational binding rules, whereas the second layer is designed to learn the deviations between these foundational scores and the actual experimental readouts under a specific experimental system.

### 2.2 Dataset Construction

#### 2.2.1 First-Layer Foundational Dataset

The positive samples for the first layer were sourced from miRTarBase Release 10.0. The inclusion criteria required the species to be *Homo sapiens* and the interactions to be supported by strong experimental evidence, such as Western blot, reporter assay, or qRT-PCR. After deduplication, 8,667 positive sample records were obtained.

The negative samples were derived from TarBase V8, filtered for the species *Homo sapiens* and a “negative” designation in the positive_negative field. After deduplication, 46,694 negative sample records were obtained, yielding a positive-to-negative sample ratio of approximately 1:5.4.

To strictly prevent data leakage, the dataset was partitioned into training, validation, and testing sets grouped by target genes. This ensured that the same target gene did not appear simultaneously in the training set and either the validation or test sets. The final partitioning results are detailed in Table 1.

**Table 1:** Partitioning statistics of the first-layer dataset.

| Subset | Total Samples | Positive Samples | Negative Samples | Negative/Positive Ratio | Number of Genes | Number of miRNAs |
| --- | --- | --- | --- | --- | --- | --- |
| Training Set | 38,752 | 6,067 | 32,685 | 5.39 | 12,453 | 820 |
| Validation Set | 5,536 | 867 | 4,669 | 5.39 | 4,268 | 379 |
| Test Set | 11,073 | 1,733 | 9,340 | 5.39 | 6,925 | 508 |
| Total | 55,361 | 8,667 | 46,694 | — | — | — |

#### 2.2.2 Second-Layer Wet-Lab Experimental Dataset

The second-layer samples are derived from the real experimental validation of endogenous expression regulation between plant-derived small RNAs and human target gene 3’UTRs (determined by significant changes in endogenous target protein expression detected by Western blot or ELISA assays 48 hours post-transfection of small RNAs via Lipo3000). The dataset comprises a total of 40 samples, consisting of 13 positive samples and 27 negative samples.

Experimental assays were conducted in cell models such as Hep3B, H9c2, corpus cavernosum cells, and UMR-106 cells. The experimental protocol involved a standard transfection for 4 hours, followed by continued incubation for 48 hours. The changes in endogenous target protein expression levels were subsequently detected by Western blot/ELISA to determine the experimental readouts for the candidate small RNA-target gene 3’UTR combinations.

Since the wet-lab experimental data provided only small RNA sequences and target gene names, we mapped the gene names to their corresponding transcripts and 3’UTR sequences using the human transcript 3’UTR sequence reference set (GRCh38) exported via Ensembl BioMart.

### 2.3 Interpretable Feature Engineering

The final feature engineering system for the first layer encompasses 41 dimensions, covering core categories such as the intrinsic sequence properties of small RNAs, global 3’UTR context, canonical site searches and site statistics, local AU/GC context, and compensatory pairing and thermodynamic features. All features were derived computationally from the small RNA sequences, candidate 3’UTR sequences, site-search results, or structural thermodynamic algorithms. Thermodynamic features were computed using the ViennaRNA Package (RNAfold v2.5.1) and RNAhybrid (v2.1.2).

The main feature categories are described below:

For samples that completely lack a canonical site, the model adopts a candidate site supplementation strategy: a sliding window is utilized to scan the 3’UTR for candidate regions with the highest number of seed matches, thereby generating comparable site descriptors. Although this strategy prevents samples without canonical sites from being excluded from the model, it may introduce a representation bias centered heavily around seed rules; this is further addressed as a limitation in the Discussion section.

The second layer ultimately utilizes 52-dimensional computational features, which encompass the sequence, site, context, and thermodynamic features corresponding to the optimal transcript, the optimal transcript score from the first layer (best_score), and the total number of candidate transcripts (transcript_count). The six thermodynamic-related fields in the second layer were all derived from actual algorithmic computations and exhibit inter-sample variance across the 40 experimental samples.

### 2.4 Model Training

#### 2.4.1 First-Layer XGBoost

The first-layer model is based on XGBoost v3.2.0, with the objective set to binary logistic regression(binary:logistic), log loss(logloss) and AUC as evaluation metrics. To balance the classes during training, we set the scale_pos_weight parameter according to the ratio of positive to negative samples in the training set. The important hyperparameters are summarized below:

The aforementioned hyperparameters were determined via an Optuna-based Bayesian search optimized against the validation set AUC. The search space encompassed parameters such as max_depth ∈ [4, 12], learning_rate ∈ [0.01, 0.1], and n_estimators ∈ [200, 1000], with the optimal combination selected after 100 trial rounds. The decision threshold was chosen by sweeping candidate thresholds across the validation set, prioritizing a recall of no less than 0.85, and then selecting the threshold that optimized precision and accuracy. The final selected threshold was 0.4797. Because the decision threshold was selected based on the validation set, the validation performance reported in Table 2 may exhibit a slight overestimation; however, the test set results represent a completely independent evaluation.

**Table 2:**
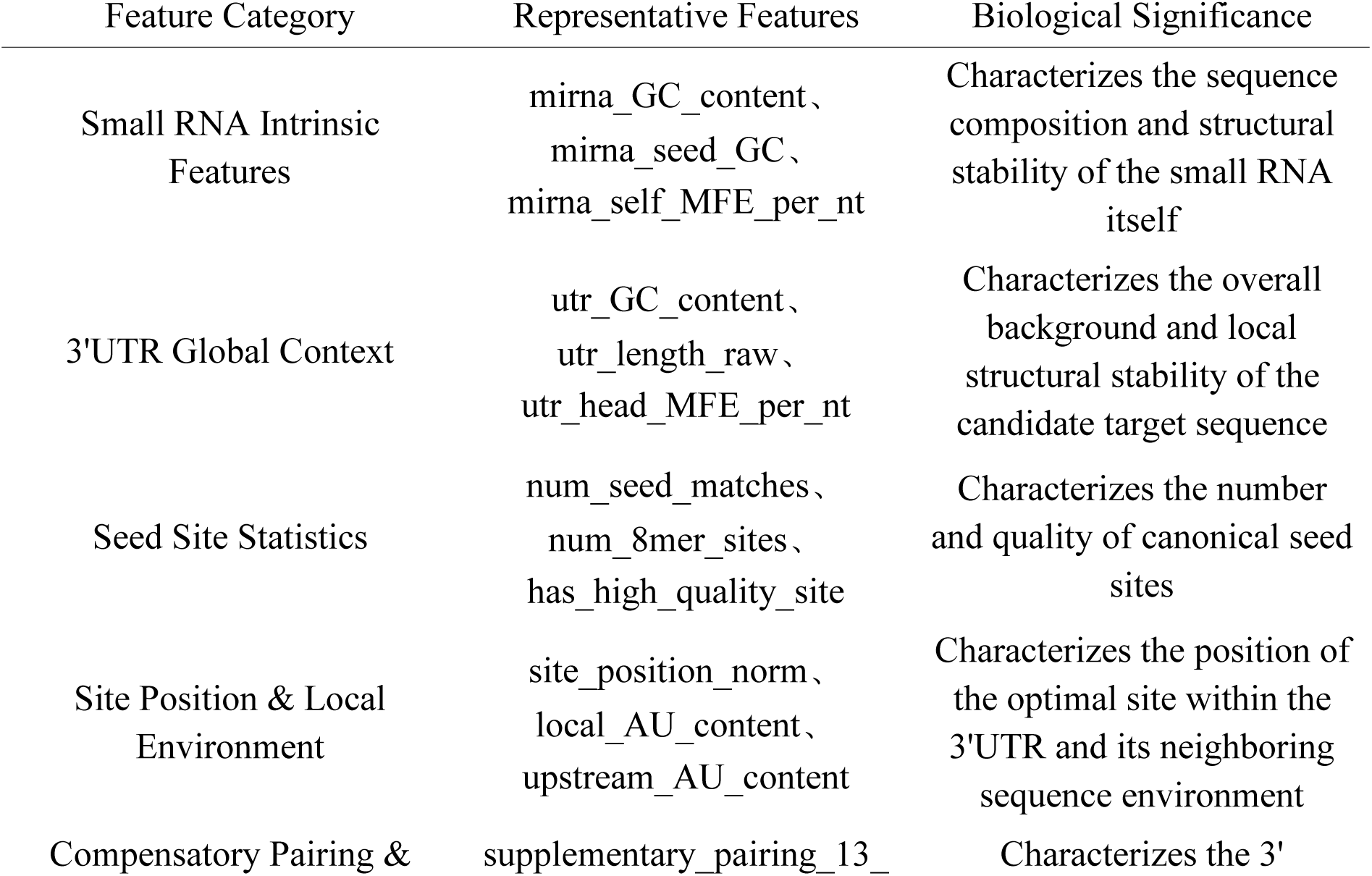
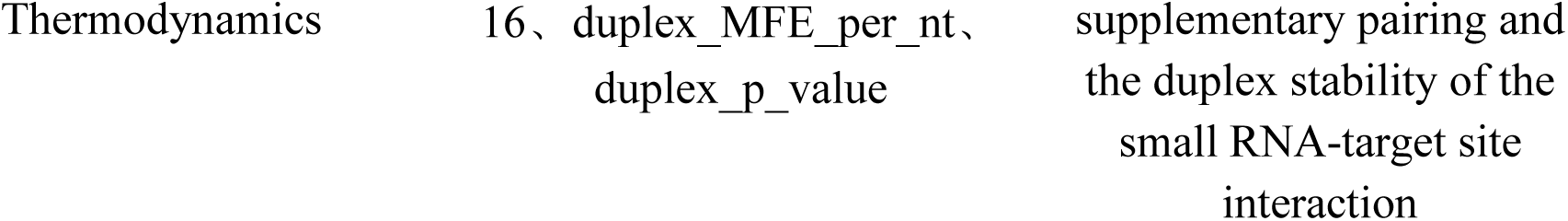
Primary interpretable feature categories and their biological significance.

**Table 3:** Critical hyperparameters of the first-layer XGBoost model.

| Parameter | Value |
| --- | --- |
| n_estimators | 500 |
| max_depth | 8 |
| learning_rate | 0.05 |
| subsample | 0.60 |
| colsample_bytree | 0.70 |
| min_child_weight | 15 |
| gamma | 0.2 |
| reg_lambda | 2.0 |
| reg_alpha | 0.1 |
| random_state | 42 |

#### 2.4.2 Second-Layer TabPFN

The second-layer model utilizes TabPFN (v0.1.11, with N_ensemble_configurations = 16 and GPU acceleration). TabPFN was selected because the second-layer sample size consists of only 40 records, making traditional high-capacity models highly susceptible to overfitting. Conversely, TabPFN possesses strong pre-trained priors tailored for small tabular classification tasks.

The performance of the second layer was evaluated using the following strategies:

1. **A single 70/30 stratified random split**, reporting test set results exclusively.
2. **Ten repeated 70/30 stratified random splits**, reporting the mean, standard deviation, and range of metrics.
3. **A 5-fold cross-validation**, serving as a complementary evaluation for the small-sample regime

For all second-layer evaluations, a first-layer baseline was simultaneously calculated by directly utilizing the first-layer best_score as the predicted probability.

### 2.5 Evaluation Metrics and Interpretability Methods

The main evaluation metrics are Area Under the Receiver Operating Characteristic curve(AUROC/AUC),Precision-Recall AUC(PR-AUC), Recall, Precision and Accuracy. For the first layer, we carried out complementary experiments, namely, gene-based group splitting,miRNA-based group splitting and label-shuffling, to rigorously evaluate the cross-entity generalization capability and avoid information leakage. We used SHAP(SHapley Additive exPlanations) values to provide model interpretability, together with feature Gain analysis to assess the feature contributions in the first-layer XGBoost model.

## 3 Results

### 3.1 The First-Layer Model Learns Stable Foundational Binding Rules

The first-layer XGBoost model achieved strong performance on the randomly split test set (Table 4).

**Table 4:** Performance of the first-layer XGBoost model under random splitting (decision threshold = 0.4797)

| Data Subset | AUC | Recall | Precision | Accuracy |
| --- | --- | --- | --- | --- |
| Training Set | 0.9994 | 0.9984 | 0.9427 | 0.9901 |
| Validation Set | 0.9700 | 0.8524 | 0.8026 | 0.9445 |
| Test Set | **0.9686** | **0.8523** | **0.8080** | **0.9452** |

From the test set metrics, we see that the XGBoost model driven by the 41-dimensional interpretable features has a high capacity to discriminate between positive and negative binding samples. The gap between the train and test sets(AUC 0.9994 vs. 0.9686) suggests a moderate degree of overfitting to the training distribution, which is our main motivation for the following group-splitting and label-shuffling experiments.

### 3.2 Generalization Boundaries: Unseen Genes Perform Better Than Unseen miRNAs

The results of the group-splitting experiments are presented in Table 5.

**Figure 2:**
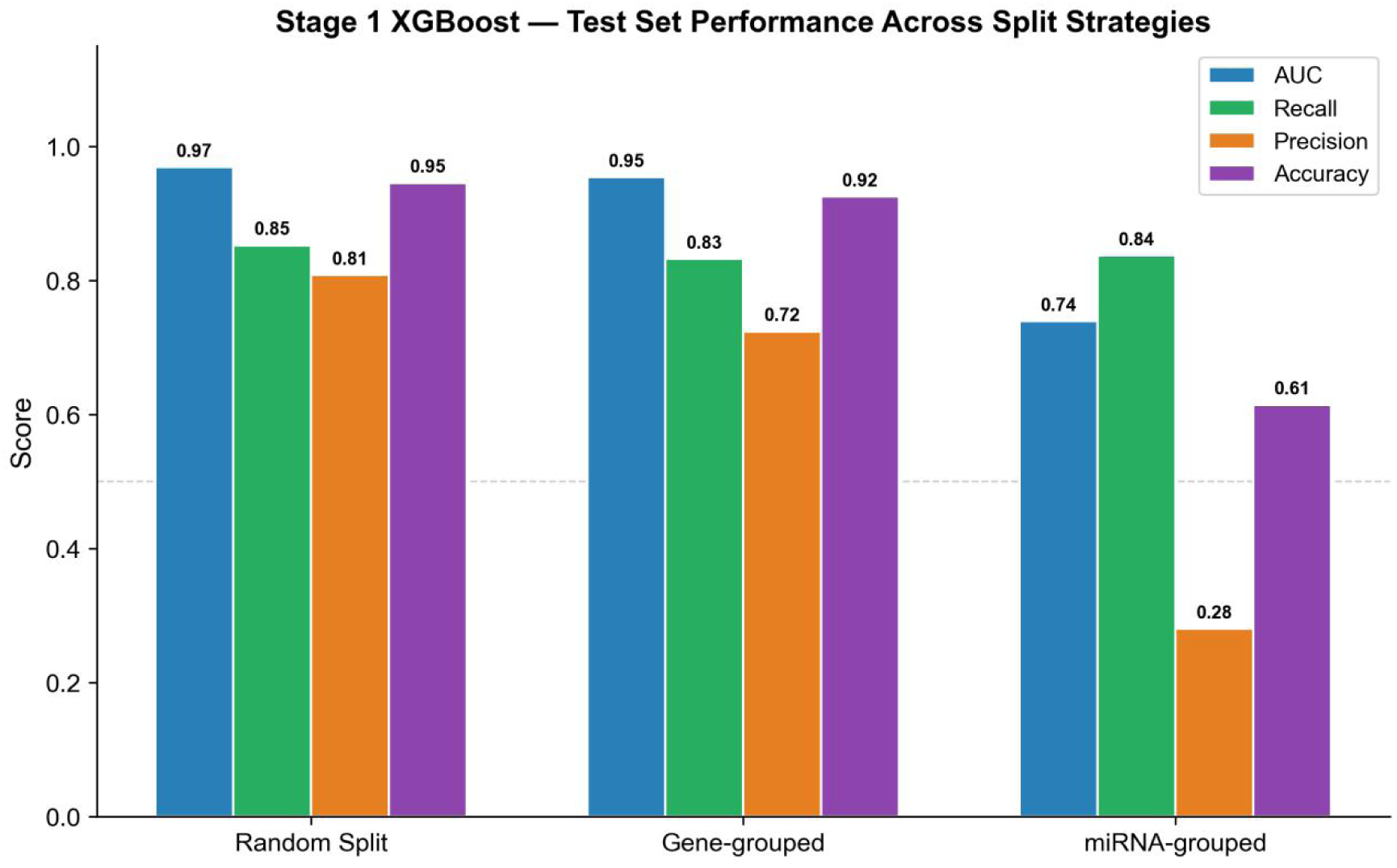
Performance comparison on the test set under three partitioning strategies.

**Table 5:** Performance comparison across different group-splitting experiments (Test Set)

| Partitioning Strategy | Decision Threshold | AUC | Recall | Precision | Accuracy |
| --- | --- | --- | --- | --- | --- |
| Random Split | 0.4797 | 0.9686 | 0.8523 | 0.8080 | 0.9452 |
| Gene-based Grouping | 0.4397 | **0.9541** | 0.8320 | 0.7236 | 0.9248 |
| miRNA-based Grouping | 0.7079 | **0.7390** | 0.8368 | 0.2799 | 0.6142 |

When grouping by gene, the model AUC is still high at 0.9541, indicating that the first layer has a good cross-entity discrimination ability for unseen target genes. When grouping by miRNA, the AUC drops to 0.7390 and the precision decreases sharply to 0.2799. This shows that the model generalization ability to small RNA sequences completely unseen by the training set is poor, a fact very consistent with the high feature contributions of small RNA intrinsic sequence properties in the SHAP analysis.

In the label-shuffling experiment, the AUC values for the training, validation, and test sets were 0.5075, 0.4900, and 0.4881, respectively. All three values hover around the random baseline of 0.5, indicating that there is no leak of labels or sample identifiers into the model inputs.

### 3.3 The Second-Layer Model Improves Adaptation Performance in Small-Sample Experiments

On the 12 test samples from a single 70/30 split, the second-layer TabPFN demonstrated a higher AUC and accuracy compared to the first-layer baseline (Table 6). This single-split result is intended solely to showcase the observed trend and should not be used as a standalone final conclusion. Across 10 repeated 70/30 random splits, the second-layer model achieved an average AUC of 0.7406 ± 0.092, which is significantly higher than the first-layer baseline of 0.3563 ± 0.149 (Table 7). Notably, the below-chance AUC of the first-layer score (0.3563 < 0.5) may suggest a weak inverse relationship between the foundational binding probability and the experimental regulatory labels in this small dataset; however, because this estimate is based on only 40 samples, it should not be over-interpreted as a robust anti-correlation, and is most plausibly explained by the distributional shift between large-scale in silico miRNA–mRNA training labels and plant-derived sRNA transfection outcomes—which motivates the use of a dedicated second-layer adapter rather than relying on the first-layer score directly. This result is consistent with the second-layer model capturing adaptation signals relevant to the experimental readouts, although it must be interpreted with caution given the limited sample size. However, because the dataset is limited to only 40 samples, a stable generalization capability toward completely novel genes or small RNAs cannot be inferred from random splits alone; we therefore additionally performed leave-one-gene-out and leave-one-small-RNA-out evaluations (Section 3.6).

**Figure 3:**
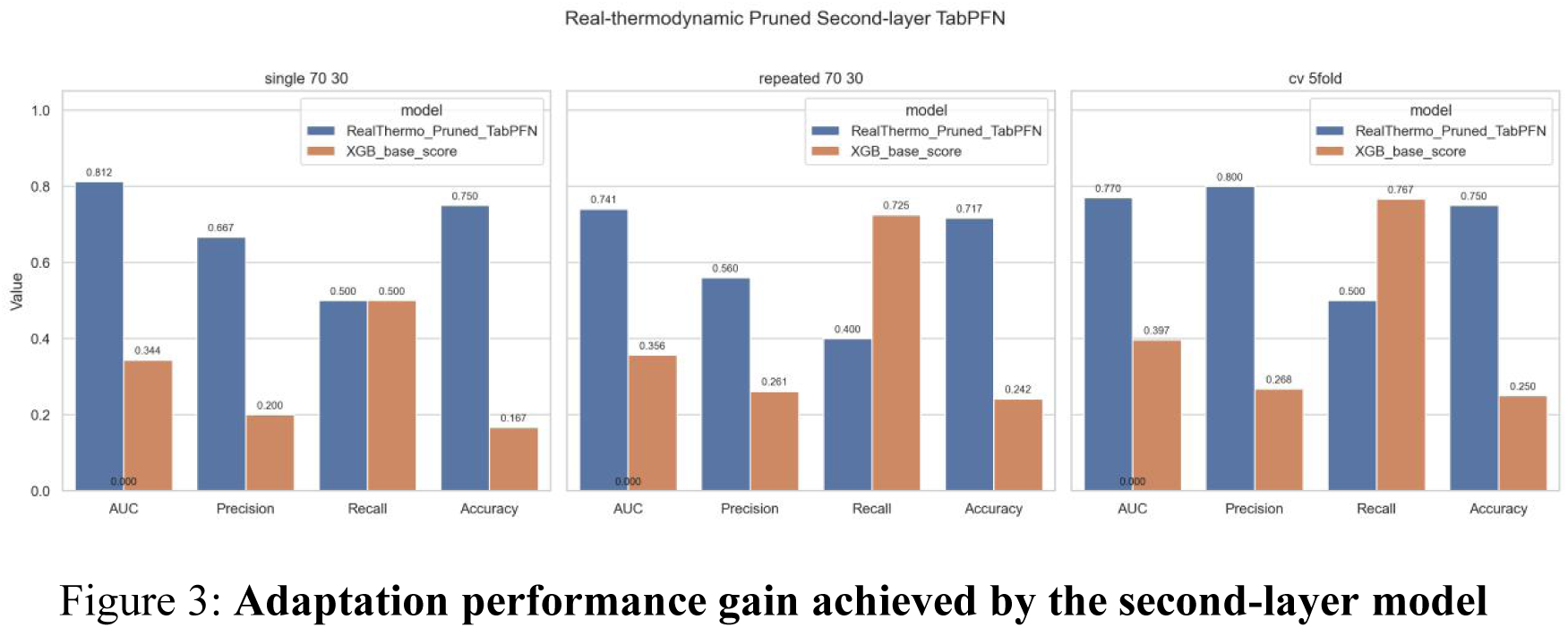
Adaptation performance gain achieved by the second-layer model.

**Table 6:**
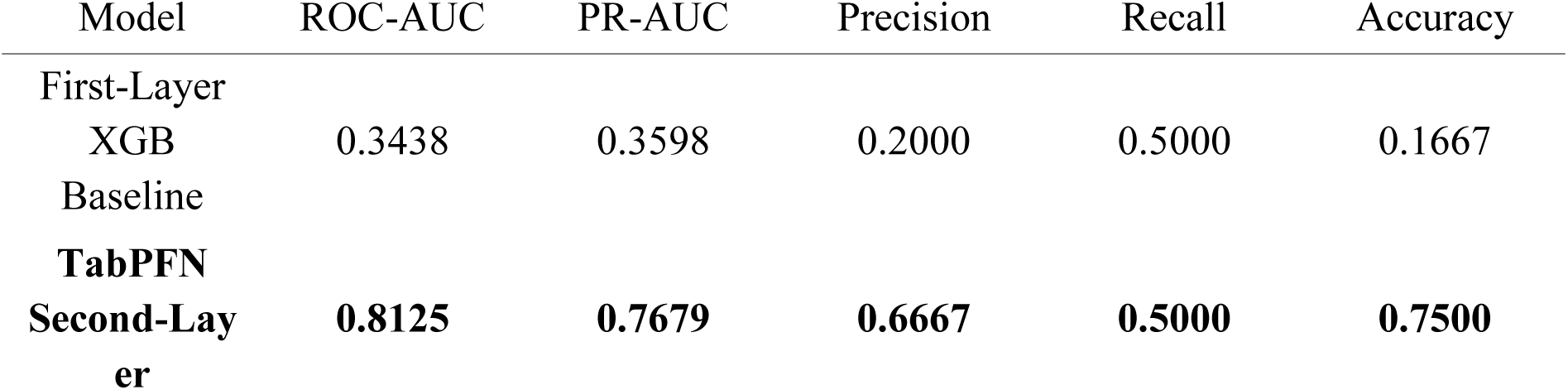
Second-layer results for a single 70/30 split test set.

**Table 7:**
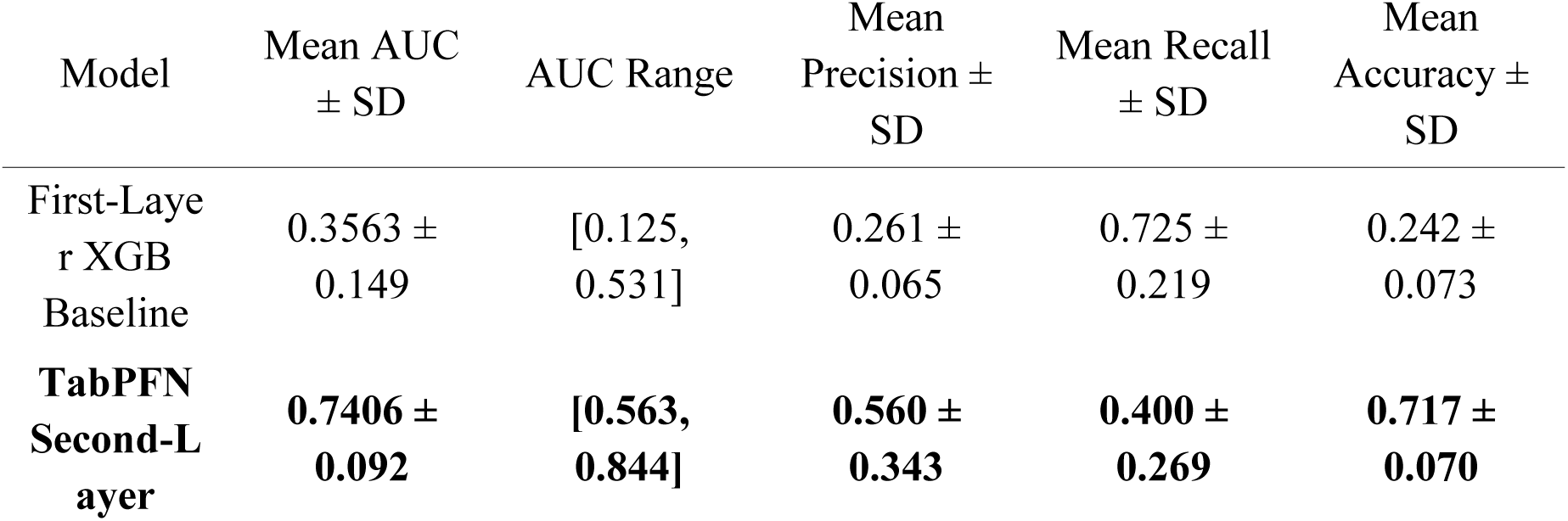
Performance on the test set across 10 repeated 70/30 splits.

In the 5-fold cross-validation, the second-layer model attained a mean AUC of 0.770±0.177, which is superior to the first-layer baseline of 0.397±0.250. Since the second layer only has 40 total samples with as few as 13 positive samples, the precision and recall values fluctuate significantly. The performance of the second-layer model should be viewed as a preliminary observation of experimental adaptation mechanisms rather than a finalized, externally validated conclusion.

### 3.4 SHAP Analysis Reveals Distinct Focus Areas Between the Two Layers

The SHAP analysis for the first-layer XGBoost model is presented in Figure 4, and the comparison between Gain and SHAP values is shown in Figure 5.

**Figure 4:**
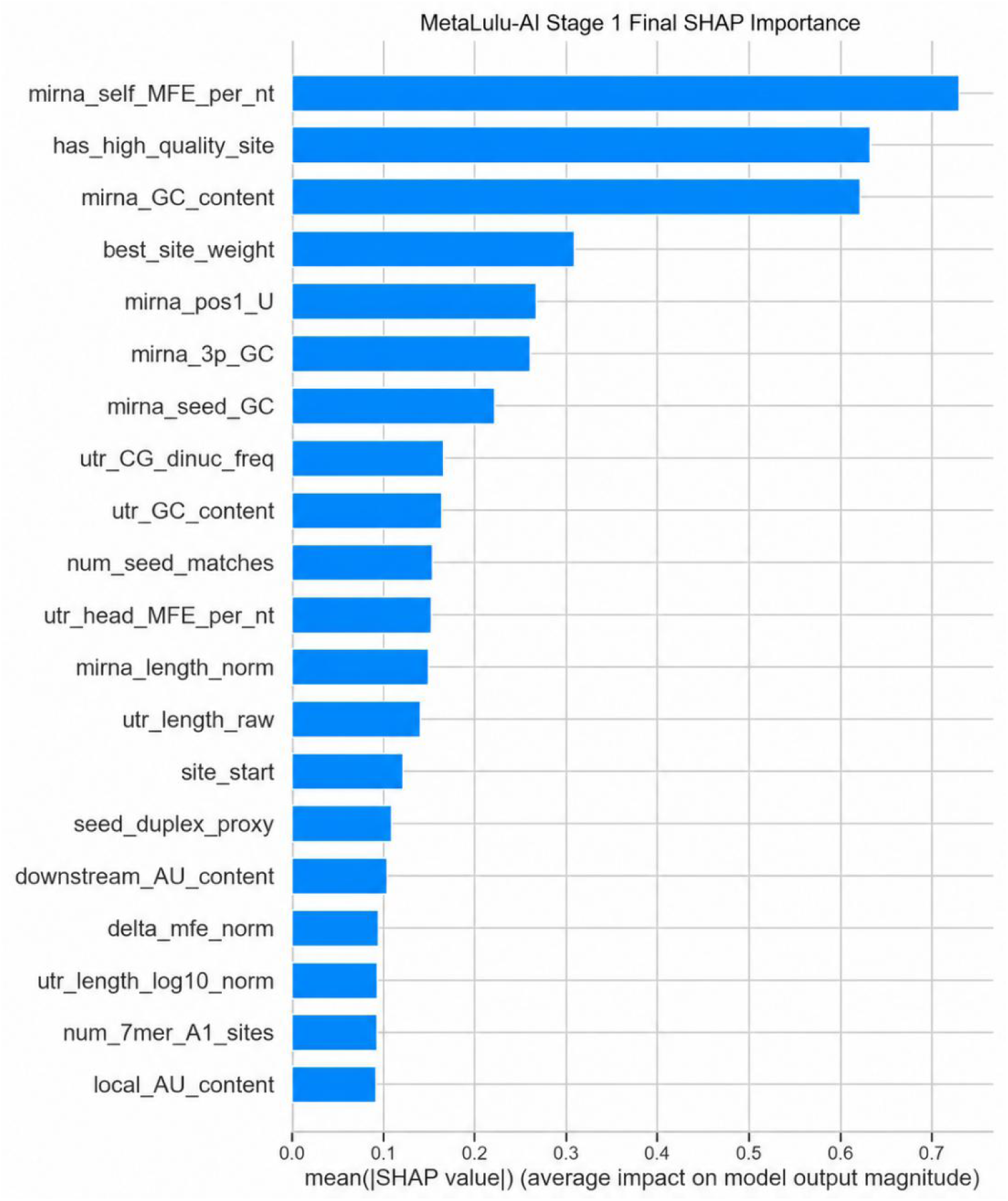
Top 20 SHAP feature importance ranking for the first-layer XGBoost model.

**Figure 5:**
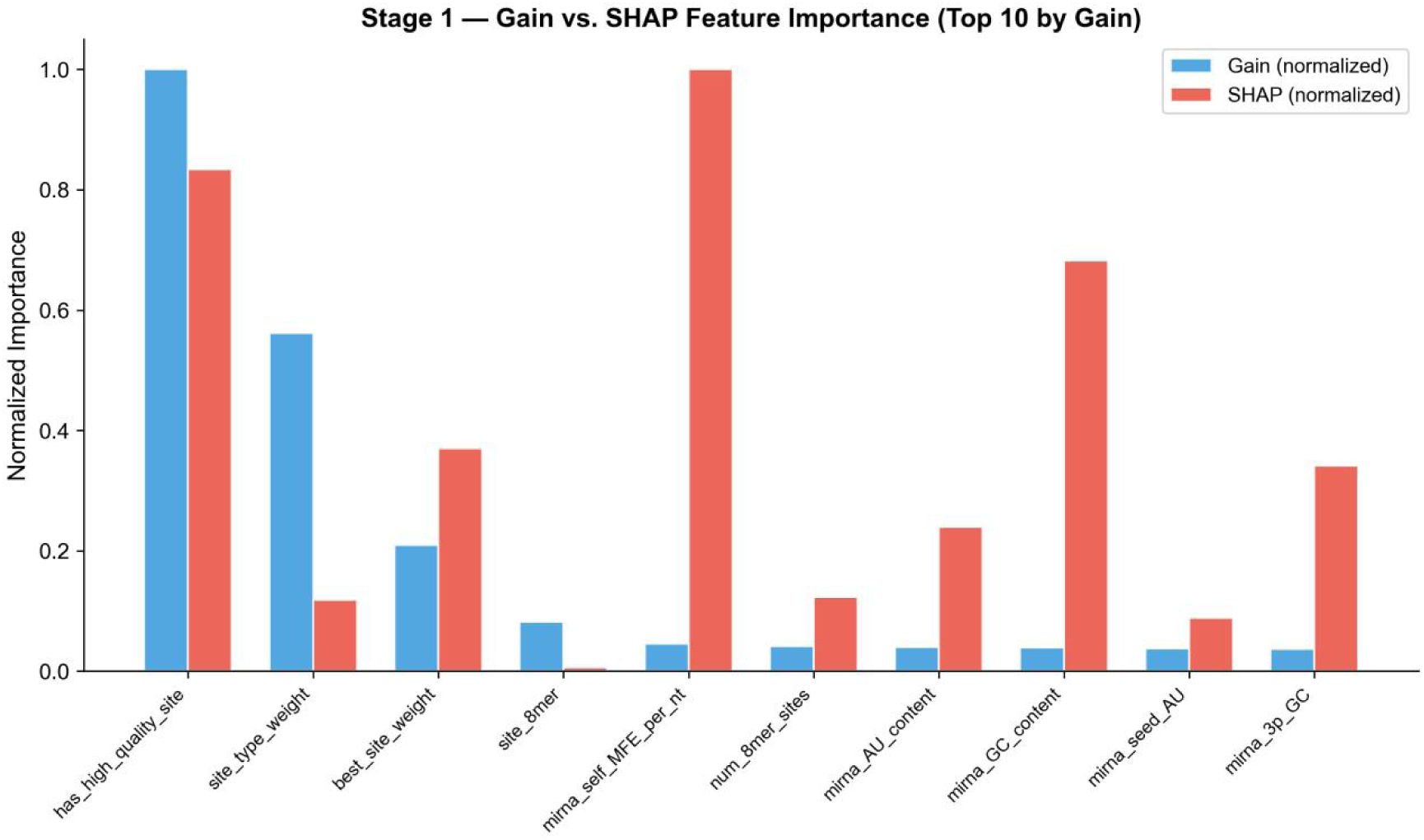
Normalized importance comparison between Gain and SHAP for the first layer (Top 10 by Gain)

The top ten features for the first layer identified by SHAP are mirna_self_MFE_per_nt, has_high_quality_site, mirna_GC_content, best_site_weight, mirna_pos1_U, mirna_3p_GC, mirna_seed_GC, utr_CG_dinuc_freq, utr_GC_content, and num_8mer_sites. This demonstrates that the first-layer model primarily leverages the structural thermodynamics of the small RNA itself, the quality of canonical seed sites, and sequence composition features to make predictions.

The SHAP analysis for the second-layer TabPFN model is shown in Figure 6.

**Figure 6:**
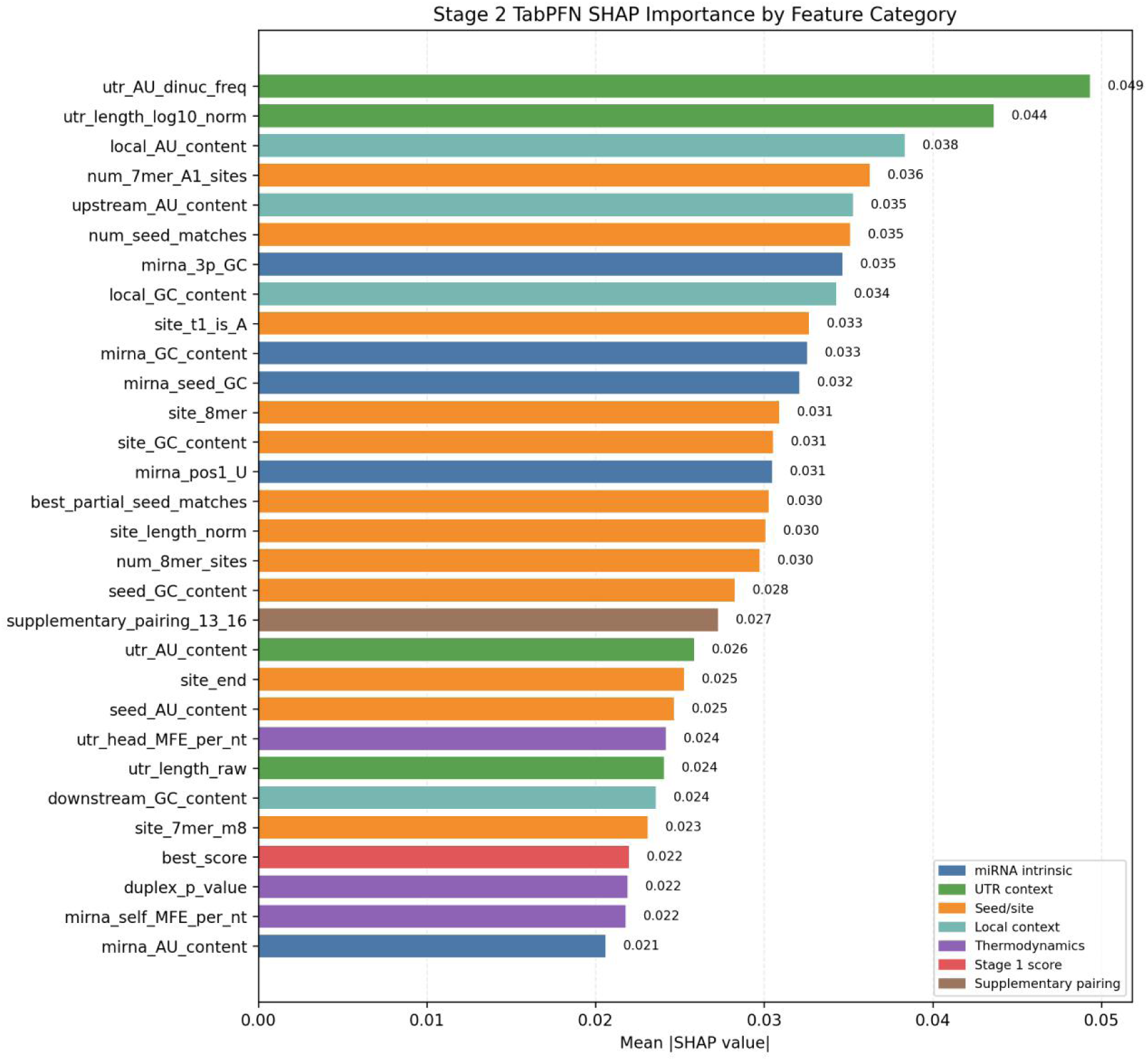
Top 30 SHAP feature importance ranking for the second-layer TabPFN model.

The top features contributing to the second layer are primarily concentrated in categories such as UTR sequence composition, site-neighboring context, and seed site statistics. The top ten features are utr_AU_dinuc_freq, utr_length_log10_norm, local_AU_content, num_7mer_A1_sites, upstream_AU_content, num_seed_matches, mirna_3p_GC, local_GC_content, site_t1_is_A, and mirna_GC_content.

Furthermore, utr_head_MFE_per_nt, duplex_p_value, and mirna_self_MFE_per_nt all rank within the top 30 of the second layer, indicating that the recomputed thermodynamic information provides an observable contribution to the experimental adaptation model. The first-layer best transcript score (best_score) also ranks within the top 30 (mean |SHAP| = 0.0220), demonstrating that while the second layer explicitly utilizes the foundational score from the first layer, its final judgment is not limited to this score alone.

### 3.5 Comparison with Classical Target-Prediction Approaches

To position the first-layer model relative to established heuristics, we benchmarked it on the held-out test set against three feature proxies of widely used tools: a seed-match score (TargetScan-style), a duplex minimum-free-energy score (RNAhybrid-style), and a combined seed-plus-MFE score (miRanda-style). As summarized in Figure 7 and Table 8, the first-layer XGBoost model substantially outperformed all three proxies, raising the AUC from the 0.656–0.808 range to 0.969 and the PR-AUC from 0.26–0.46 to 0.914. These proxies are simplified reimplementations of the corresponding scoring principles rather than the full published pipelines, so the comparison should be read as indicative of the gain from combining 41 interpretable features within a learned model rather than as a head-to-head benchmark against the released tools.

**Figure 7:**
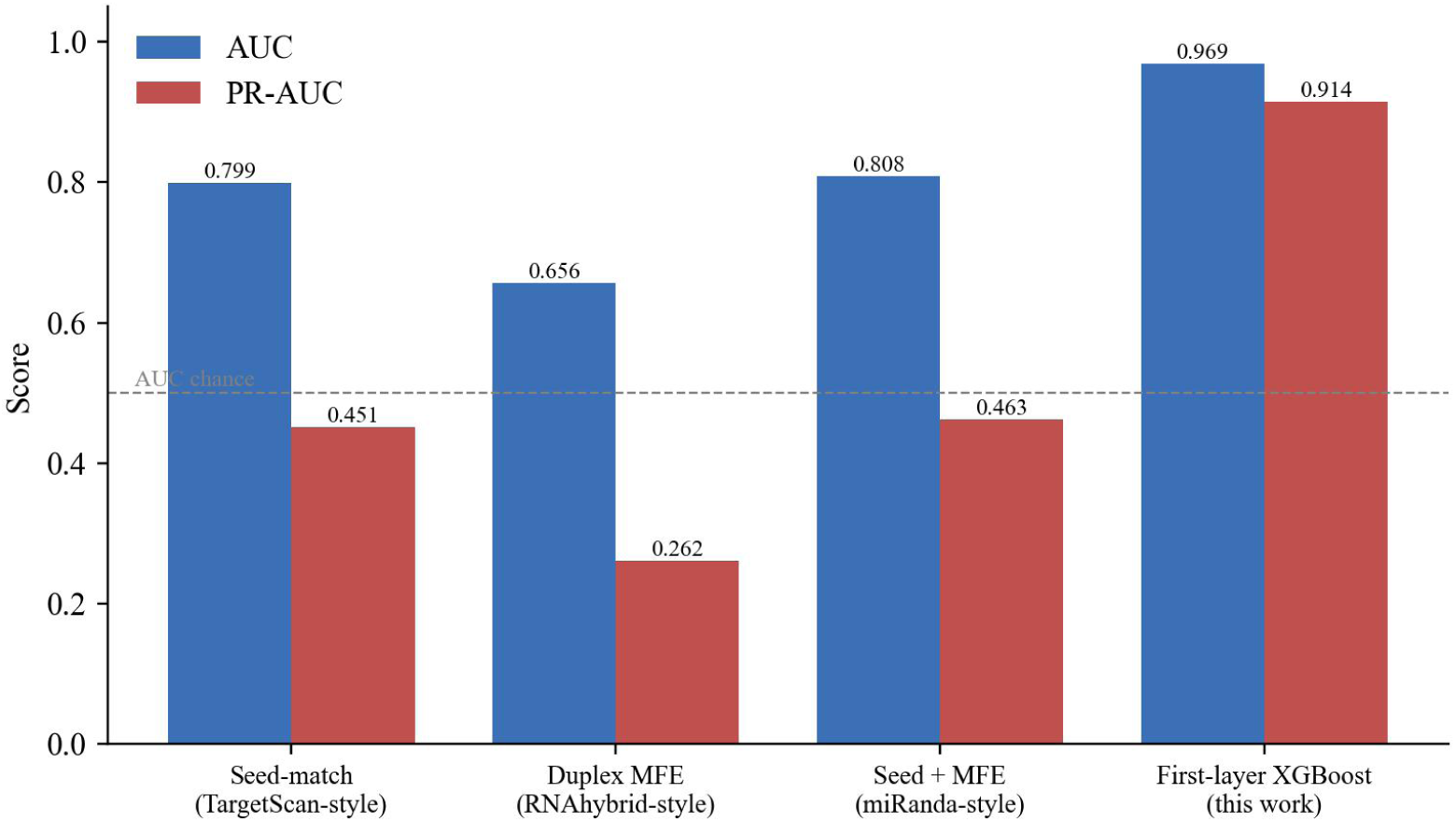
First-layer model versus simplified proxies of classical target-prediction tools on the held-out test set (AUC and PR-AUC)

**Table 8:** First-layer performance versus simplified proxies of classical target-prediction tools (held-out test set)

| Method | AUC | PR-AUC | Best F1 | Precision | Recall | Brier |
| --- | --- | --- | --- | --- | --- | --- |
| Seed-match<br>(TargetScan-style proxy) | 0.799 | 0.451 | 0.529 | 0.462 | 0.618 | 0.153 |
| Duplex MFE<br>(RNAhybrid-style proxy) | 0.656 | 0.262 | 0.330 | 0.220 | 0.659 | 0.195 |
| Seed + MFE<br>(miRanda-style proxy) | 0.808 | 0.463 | 0.535 | 0.462 | 0.635 | 0.140 |
| First-layer XGBoost<br>(this work) | 0.969 | 0.914 | 0.830 | 0.808 | 0.852 | 0.042 |

### 3.6 Leave-One-Gene-Out and Leave-One-Small-RNA-Out Evaluation

Because random 70/30 splits can place the same gene or small RNA on both sides of the partition, we additionally evaluated the second layer under two strict entity-held-out protocols: leave-one-gene-out (LOGO), in which all samples of one target gene are held out in turn, and leave-one-small-RNA-out (LOSO), in which each unique small RNA is held out in turn. In both settings the TabPFN adapter was compared against the first-layer score and against the trivial majority-class (all-negative) baseline; because the dataset contains 13 positive and 27 negative samples, the majority-class accuracy is 0.675.

Under LOGO, the adapter reached a pooled accuracy of 0.675, far above the first-layer score baseline (0.250) but identical to the majority-class baseline. Performance was strongly gene-dependent: it was high on the class-imbalanced genes CD38 (0.90; 1/10 positive) and PDE5A (0.82; 2/11 positive), but only at chance for the balanced gene EGLN1 (0.50; 5/10 positive) and below chance for SOST (0.44; 5/9 positive). Under LOSO, the adapter reached a pooled accuracy of 0.725, marginally above the majority baseline, by correctly classifying 24 of 27 negative small RNAs (0.89) but only 5 of 13 positive small RNAs (0.385). Together (Figure 8 and Table 9), these results indicate that while the second layer clearly improves on the raw first-layer score, it does not yet demonstrate robust generalization to entirely unseen genes or small RNAs—particularly for the positive (regulatory) class—and should therefore be regarded strictly as a proof of concept pending a larger experimental dataset.

**Figure 8:**
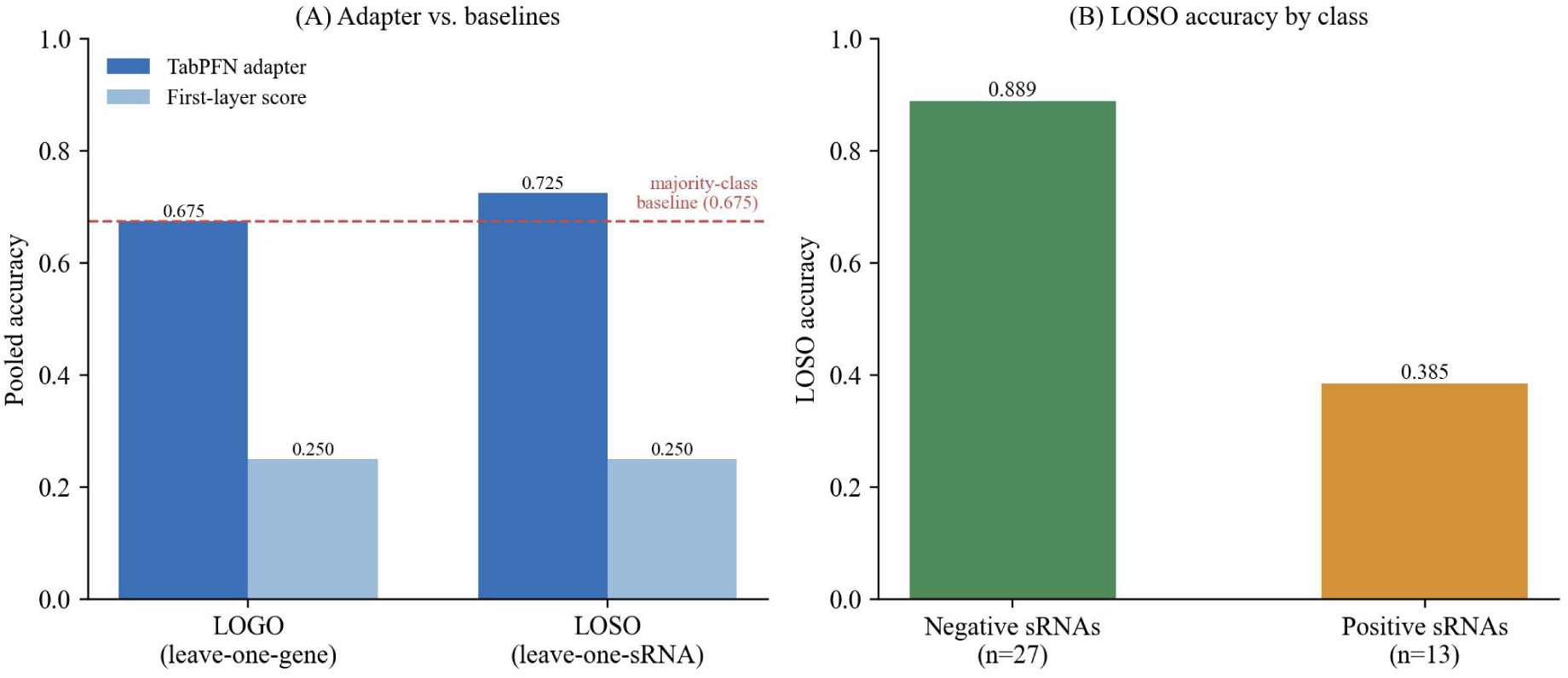
Entity-held-out evaluation of the second layer. (A) Pooled accuracy under leave-one-gene-out (LOGO) and leave-one-small-RNA-out (LOSO), with the majority-class baseline (0.675) shown as a dashed line (B) LOSO accuracy split by class, showing recovery of negatives but weak performance on positives.

**Table 9:** Second-layer accuracy under leave-one-gene-out and leave-one-small-RNA-out evaluation. Positive-class recall is not separable per fold for LOGO.

| Evaluation | TabPFN adapter acc. | First-layer score acc. | Majority-class acc. | Positive-class recall |
| --- | --- | --- | --- | --- |
| Leave-one-gene-out (4 genes, 40 samples) | 0.675 | 0.250 | 0.675 | — |
| Leave-one-small-RNA-out (40 small RNAs) | 0.725 | 0.250 | 0.675 | 0.385 (5/13) |

## 4 Discussion

### 4.1 The Necessity of the Two-Layer Architecture

Rather than employing a single end-to-end model to forecast experimental readouts directly from raw sequences, this study adopts a two-layer architecture, primarily to address the severe asymmetry in data scale. While tens of thousands of data points are available for public human miRNA-mRNA interactions, our wet-lab experimental dataset for plant-derived small RNAs consists of only 40 samples. Attempting to train a comprehensive end-to-end model—encompassing everything from sequence feature extraction to experimental readout prediction—on just 40 samples would inevitably lead to severe overfitting. The essence of this two-layer architecture is a transfer learning strategy: the large-scale dataset is leveraged to handle feature representation and foundational discrimination, whereas the small-sample dataset is dedicated solely to mapping the final step from foundational scores to experimental labels. In scenarios where the sample size is extremely constrained, this division of labor proves substantially more robust than an end-to-end approach.

The first-layer model was able to maintain a high AUC when splitting by gene, indicating that it was able to generalize to unseen target genes. The drop in performance when splitting by miRNA indicates that the model was relying too heavily on the sequence composition and intrinsic structural features of the small RNAs. This is especially important for the application of plant-derived small RNAs, since cross-kingdom plant-derived small RNAs may have different distributional differences than human endogenous miRNAs.

### 4.2 Rational Interpretation of the Second-Layer Results

The second-layer model consistently outperformed the first-layer baseline in both repeated random splits and 5-fold cross-validation, indicating the presence of learnable adaptation signals within the small-sample experimental data. However, because the dataset is limited to only 40 samples, the current performance metrics cannot support robust conclusions regarding strong generalization. We additionally performed leave-one-gene-out and leave-one-small-RNA-out evaluations for this layer (Section 3.6); under these strict entity-held-out settings the adapter still far exceeded the first-layer score baseline (pooled accuracy 0.675 and 0.725 versus 0.250) but only matched or marginally exceeded the majority-class baseline (0.675) and recovered few positive cases for unseen genes and small RNAs, so robust entity-level generalization remains unproven. Consequently, these second-layer results should be strictly interpreted as a proof of concept and a foundational baseline to guide and justify future large-scale experimental data collection.

### 4.3 Limitations

This framework is subject to several key limitations that warrant explicit discussion. The most direct constraint stems from the data scale of the second layer: a dataset of only 40 samples lacks the breadth needed to support robust generalization conclusions across diverse genes or small RNAs. Although leave-one-gene-out and leave-one-small-RNA-out evaluations were performed (Section 3.6), they showed that the adapter does not yet reliably exceed a majority-class baseline for entirely unseen genes or small RNAs, particularly on positive (regulatory) cases; the performance of the second-layer model must therefore strictly be interpreted as a proof-of-concept level observation.

For the first-layer model, the drop in AUC to 0.7390 during miRNA-based group splitting reveals a structural vulnerability: the model exhibits insufficient generalization capacity toward small RNA sequences entirely absent from the training set. Given that the sequence space of plant-derived small RNAs systematically differs from that of human endogenous miRNAs, this bottleneck could be substantially more pronounced in practical deployment than the random-split performance metrics might otherwise imply.

Regarding the feature engineering system, the current 41-dimensional feature set relies heavily on the canonical seed match hypothesis. Non-canonical regulatory modalities—such as G:U wobble base-pairing, centered sites, and 3’-compensatory-dominant binding—are only partially accounted for by existing features (e.g., supplementary_pairing_13_16) and are not yet systematically captured by the framework.

At the experimental platform level, the training and validation data utilized for the second layer were generated exclusively from endogenous protein expression assay systems (via Western blot or ELISA) following small RNA transfection. Although these data span multiple cellular environments, including Hep3B and H9c2 cell lines, we have not evaluated whether alternative modalities, such as qRT-PCR data, would yield comparable performance within the same modeling architecture. Furthermore, the negative samples for the first-layer model were sourced from the negative annotations in TarBase v8. These records inherently denote “no regulatory effect detected under specific experimental conditions” rather than a definitive “confirmed absence of physical binding,” a distinction that may introduce systematic label noise. Finally, and most fundamentally: this model serves strictly as a computational candidate-prioritization tool; it cannot substitute for systematic in vivo experimental validations concerning the absorption, stability, and actual physiological effects of plant-derived small RNAs.

### 4.4 Future Research Directions

Future work will center around establishing a closed-loop active learning framework (Figure 9). First, high-confidence predicted targets will be selected for systematic wet-lab experimental validation. Subsequently, validated positive samples will be integrated into the training set, while validated negative samples will serve as high-quality true negatives to refine the model’s decision boundaries. Finally, feature contributions and thresholding strategies will be dynamically re-evaluated based on the newly augmented dataset.

**Figure 9:**
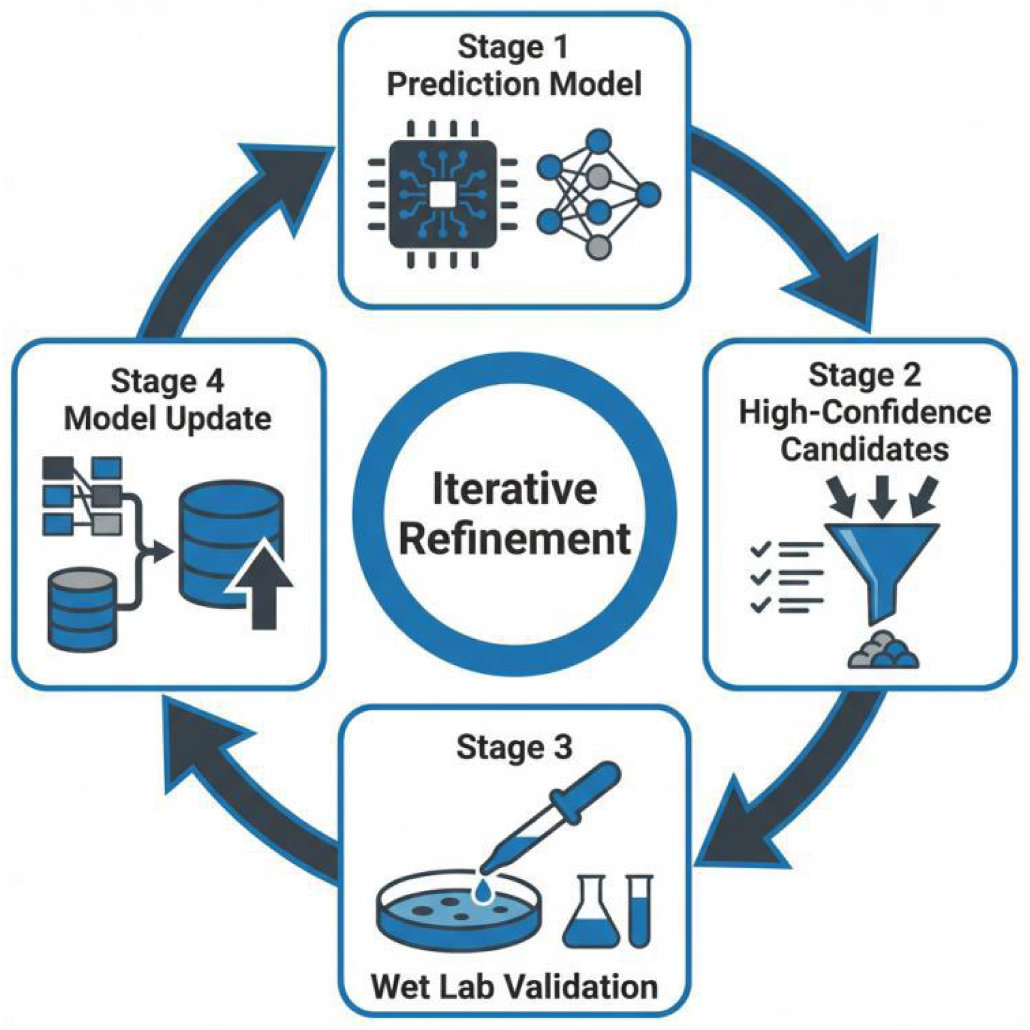
Schematic diagram of the closed-loop active learning framework.

As experimental data accumulates, the leave-one-gene-out and leave-one-small-RNA-out evaluations reported here (Section 3.6) can be repeated at larger scale and extended with cross-cell-line validation, providing a more reliable assessment of practical value in screening cross-kingdom targets for plant-derived small RNAs. Having already benchmarked the first layer against seed-match, duplex-MFE, and combined proxies of TargetScan, RNAhybrid, and miRanda (Section 3.5), future efforts will extend these comparisons to the full published tools and to simple linear classifiers to further clarify the source of performance gains and the operational boundaries of the TabPFN adapter.

## 5 Conclusion

This study comprehensively reports the design architecture and preliminary evaluation results of the proposed framework. The framework breaks down the prediction of cross-kingdom plant-derived small RNAs targeting human mRNA 3’UTRs into a two-stage process: the first layer learns foundational binding principles from 55,361 public human miRNA-mRNA data points, achieving a test set AUC of 0.9686 and maintaining an AUC of 0.9541 during gene-based group splitting; the second layer performs system adaptation on 40 experimental data points of endogenous protein expression regulation (via Western blot/ELISA), achieving an average AUC of 0.7406 across 10 repeated splits—marking a substantial improvement over the direct utilization of first-layer scores (0.3563). A comparison of the SHAP analyses between the two layers reveals a divergence in feature attribution: foundational binding discrimination relies more heavily on the intrinsic characteristics of the small RNA, whereas experimental adaptation is predominantly driven by the local UTR genomic context.

Although the proposed framework shows preliminary potential for cross-kingdom regulatory prediction, we maintain a rigorous and cautious scientific perspective regarding the experimental adaptation results at this current stage. A proof of concept based on 40 samples remains far from a robust, generalized conclusion, and our leave-one-gene-out and leave-one-small-RNA-out tests confirm that generalization to unseen genes and small RNAs is not yet established. The core objective of future work will be to expand experimental data coverage through a closed-loop active learning framework and to validate the transferability of this framework across a wider array of target genes and experimental platforms.

## Data and Code Availability

All data and code to reproduce the analyses in this manuscript are available at https://github.com/whiteqiao6253/metalulu-ai. The 40 wet-lab experimental samples used for Stage 2 model training are provided as stage2_experimental_data.csv in the repository. The Stage 1 training data were sourced from miRTarBase Release 10.0 (https://mirtarbase.cuhk.edu.cn/) and TarBase v8 (http://diana.imis.athena-innovation.gr/DianaTools/), both of which are publicly available databases.

## Ethics Statement

This study did not involve human participants, patient-derived samples, or animal subjects. All experiments were performed using commercially available cell lines; therefore, no ethical approval was required.

## Author Contributions

Weizhong Li conceived and designed the study and performed the experiments. Le Qiao developed the computational models and wrote the manuscript. Both authors read and approved the final manuscript.

## Supporting information

stage2 experimental data

